# Revisiting Evidence for Epigenetic Control of Alternative Splicing

**DOI:** 10.1101/2024.08.30.610315

**Authors:** Quirin Manz, Markus List

**Author notes:** Corresponding author(s). E-mail(s); Contributing authors.

## Abstract

Alternative splicing is crucial for increasing eukaryotic cell transcriptome and proteome diversity. Changes in alternative splicing play a key role in cell differentiation and tissue development, and aberrations in this process have been associated with diseases. Despite its importance, the exact mechanisms for regulating alternative splicing are poorly understood. Several epigenetic marks, such as histone modification H3K36me3, have previously been associated with changes in alternative splicing. Here, we leverage the EpiATLAS data set to systematically re-evaluate evidence for epigenetic control of alternative splicing.

We used SUPPA2 to calculate percentage spliced-in (PSI) values for skipped exons and retained introns and integrated this information with histone ChIP-seq and DNA methylation data. In addition to genome-wide association analysis with partial correlation and machine learning, we perform locus-specific PSI modeling on individual alternative splicing events. The latter represents a new contribution enabled by the unprecedented number of uniformly processed datasets in EpiATLAS.

Our results confirm previously reported global associations of DNAm and H3K36me3 for exon inclusion and emphasize the importance of intrinsic features for genome-wide associations.

On an event-specific level, trying to identify co-transcriptionally spliced events with epigenetic influence, we show that overall gene expression biases locus-specific analyses. Further, we show that epigenetic signal can predict PSI value in cis and trans, indicating sample-specific epigenetic fingerprints that distort across-sample analyses. Specifically, generalized linear models select epigenetic features predictive for PSI values not only in their genomic vicinity but also on different chromosomes.

Our study demonstrates that epigenetic marks are associated with the alternative inclusion of exons and introns. Without a mechanistic explanation for these associations, our work emphasizes the need for more detailed research into the relationship between epigenetic changes and transcriptome diversity.

## 1 Introduction

The process of alternative splicing alternative splicing (AS), in which introns are removed while exons are joined together variably, creates different transcripts for the same gene and thus is a major contributor to proteome diversity. AS is also linked to various diseases. For instance, in autism spectrum disorder (ASD), a misregulation of micro-exons leads to decreased protein interactivity [1]. Brain development and dysfunctions are also associated with AS [2]. Furthermore, AS plays a role in many cancer types [3–5]. Recently, there have been efforts to use the knowledge about splice variants in developing therapeutic approaches, e.g., for spinal muscular atrophy [6]. Splicing patterns vary widely between different cell types and tissues, suggesting the involvement of regulatory mechanisms that are currently insufficiently understood. Yet, understanding these mechanisms is a prerequisite for developing new treatment options.

Currently, two models for splicing regulation are discussed in the literature. The *kinetic coupling model* states that the elongation rate of the RNA polymerase II (Pol II) during the transcription process mediates splicing through a window of opportunity. All molecular components must be assembled, and the necessary reactions must occur in this limited time. While it was previously believed that splicing is a post-transcriptional process, most splicing occurs co-transcriptionally and may thus be influenced by the chromatin state. The *recruitment model* posits that the chromatin-adaptor complexes, composed of histone-binding proteins, recruit splicing factors to the nascent pre-messenger RNA (mRNA). This theory implies a regulatory role of changes to the histone proteins that promote or suppress the interaction with other proteins.

These two models are not mutually exclusive. Even the drivers of both models, i.e., histone marks and Pol II elongation rates, are known to influence each other [7, 8]. Another recent finding is the influence of genomic regulatory elements on splice variants [9, 10]. Regulatory elements are commonly linked to gene expression suppression or enhancement by binding chromatin remodelers or transcription factors [11]. They contribute to the dynamic nature of gene regulation and thus could also add to the complex splicing mechanism.

Principal determinants of splicing are sequence motifs that allow spliceosome components to bind to the ribonucleic acid (RNA). Depending on their exact sequence, these splice sites can be weaker or stronger and influence splicing behavior [12, 13]. Mutations are also known to disrupt or create splicing signals. Other contributors to splicing are splicing factors such as SR or hnRNP proteins [14]. They bind to short sequences on the nascent RNA that can enhance or suppress the use of a splice site [15]. These sequences are called exonic/intronic splicing enhancer/suppressor, depending on their location, i.e., within an intron or exon. RNA secondary structures like hairpin loops can hide potential splice sites or bring RNA-binding proteins in spatial proximity to each other [16]. Based on the sequence features, remarkable advances have been made in predicting AS. *SpliceAI* [17] is a deep learning tool that can successfully determine splicing outcomes from mutations in disease contexts. Building on that architecture, *Pangolin* [18] proved that tissue-specific splice site usage predictions are possible from sequence alone. However, it is unclear how to interpret such predictions as the sequence is constant for different tissues and cell types. It can thus be hypothesized that epigenetic mechanisms are involved in splicing regulation.

In this study, we first revisit the literature for evidence of epigenetic control of AS. Next, we leverage the most comprehensive set of reference epigenomes to date, the International Human Epigenome Consortium (IHEC) EpiATLAS [19, 20], to systematically assess the role of histone and DNA modifications in AS.

### 1.1 RNA Polymerase II as Driver

The connection of histone modifications to the Pol II elongation rate has been studied extensively. In the early 2000s, *Kornblihtt* and colleagues found that the acetylation of histones three and four can inhibit the inclusion of an exon in the fibronectin gene through elevated Pol II elongation rate [21, 22]. This result was later also linked to fewer free splicing factors [23]. Additionally, an increase in Pol II elongation rate along with H3K9ac and H3K36me3 are associated with excluding an alternative exon in *NCAM* [24]. Conversely, the suppressive histone marks H3K9me2 and H3K27me3, together with a decreased Pol II elongation rate, are associated with including the same exon [25]. Saint-André et al. showed that a heterochromatin protein 1 (HP1) protein associates with H3K9me3 and alternative exon inclusion in multiple genes linked to lower Pol II elongation rate [26]. HP1 proteins also recruit splicing factors in an exon-specific fashion, as investigated by Yearim et al.. While they peripherally related this mechanism to Pol II elongation, they mainly based it on H3K9me3 and, most prominently, DNA methylation (DNAm) [27]. Later, Batsché et al. found reduced H3K9me3 and HP1*γ* associated with exon exclusion [28]. Although they also made the connection to reduced Pol II elongation rates, they identified DNAm as the driver.

Binding of the CTCF protein to the deoxyribonucleic acid (DNA) can also cause Pol II pausing, thus including an exon in the CD45 gene [29]. Shukla et al. also found a converse effect for DNAm and exon inclusion since methylated cytosines prevent CTCF-binding. DNAm can also cause exon inclusion through the MeCP2 protein, which recruits HDACs and causes Pol II pausing [30].

The role of Pol II in influencing alternative exon and intron inclusion is not as straightforward as initially suggested. Fong et al. identified *class I* (following the abovementioned principle) and *class II* (showing opposing behavior) exons [31]. They also revealed that some constitutive exons show alternative splicing if the Pol II elongation rate deviates from the standard rate. However, these exons exhibit the same tendency (inclusion or exclusion) independent of the direction (increase or decrease in the elongation rate).

### 1.2 Histone Modifications Recruit Splicing Factors

Sims et al. showed that H3K4me3 can recruit CHD1 which binds the U2 small nuclear ribonucleoprotein (snRNP), thus assisting the splicing process [32].

In 2010, Luco et al. showed that low H3K27me3, H3K4me3, and H3K9me1 and high H3K36me3 and H3K4me1 were associated with exon exclusion of the first of two mutually exclusive exons (MX) in the FGFR2 gene. Specifically, these PTB-dependent exons are skipped through interaction with MRG15, promoted by H3K36me3 [33]. Segelle et al. caused the inclusion of the second of the MX in FGFR2 by increasing H3K27ac and decreasing H3K27me3. Further, they found a PTB-dependent, MRG15-independent exon in the CTNND1 gene whose inclusion could be induced by H3K27me3 increase and H3K27ac decrease [34].

H3K36me3 can also recruit serine/arginine-rich splicing factor 1 (SRSF1) through a protein isoform encoded in the PSIP1 gene. Pradeepa et al. found this mark and splicing factor enriched in expressed exons [35].

### 1.3 Genome-Wide Analyses

Identifying epigenetic mechanisms regulating particular AS events has brought important insight into the molecular machinery. Nevertheless, the experimental validation of such regulatory processes is time and cost-intensive. Genome-wide approaches, often supported by machine learning techniques, have become prevalent through advances in high-throughput sequencing and computer technologies.

Kolasinska-Zwierz et al., Hon et al., and Shindo et al. independently found H3K36me3 enriched globally in included alternative exons [36–38]. Conversely, H3K9me3 [37], H2BK12ac, H3K4me1, H3K4me2, H3K4me3, H3K9ac, and H4K5ac were depleted in these exons [38].

Enroth et al. built a rule-based model to apply the *histone code* to decipher the *splicing code* [39]. Using 38 binarized histone modifications at the preceding, alternative, and succeeding exon, they found H3K36me3 negatively associated with both included and excluded alternative exons. Interestingly, all top-ranked rules predicting inclusion required marks to be absent, while complex combinations arose for exclusion. The limitations of this study are the poor H3K36me3 quality and the application of a rule-based model to a high-complexity problem.

Further substantiating the positive association of H3K36me3 with the inclusion of alternative regions, Zhou et al. also identified other epigenetic marks linked to alternative splicing [40]. Those include H3K27me3, H3K4me3, DNAm, nucleosome occupancy, H3K4me2, H3K9me1, H3K79me2 and the proteins Pol II, EGR1, GABP, SRF, and SIN3A. It is important to note that Pol II chromatin immunoprecipitation sequencing (ChIP-seq) signal cannot be directly mapped to its elongation rate. An essential drawback of this study is that it analyzes epigenetic signals in predefined regions without comparing them to measured transcript expression.

Zhu et al. showed a positive linear relationship between H4K20me1 and H3K36me3 signals and skipped exon (SE) inclusion levels [41].

Podlaha et al. investigated twelve histone marks in nine cell lines and examined their correlation with mRNA and lincRNA splicing and transcription start site (TSS) selection, also considering regions further from the alternative exons [42]. On a transcriptome-wide level, exon inclusion in protein-coding transcripts positively correlated with H3K36me3 and negatively with H3K4me2 and H3K4me3. Additionally, their exon-specific analysis revealed significant gene-specific associations for H2az, H3K4me1, H3K4me2, H3K4me3, H3K9ac, H3K9me3, H3K27ac, H3K27me3, H3K36me3, H3K79me2, and H4K20me1. Unfortunately, the authors did not report the direction of these connections.

A study by Liu et al. addressed this direction problem by using logistic regression to predict SE inclusion [43]. Again, H3K36me3, H3K9me3, H3K27me3, and H4K20me1 were found to have a strong positive association. This relationship is reversed for the three latter marks when looking at the region succeeding the exon.

Chen et al. found predictive marks for SE inclusion and exclusion. Among the 20 most relevant features were only H2BK5ac, H3K9me2, and H3K79me2 inside the alternative exon, and all other predictors contained marks in the regions pre- or succeeding the exon [44]. In 2018, Chen et al. built multiple machine learning classifiers and compared their performance and feature selection methods. Again, most of the 81 optimal features were in the neighboring regions of the alternative exon, with only 20 inside of it. Interestingly, H3K36me3 showed a bias towards exclusion [45]. To unveil the conditional dependencies of histone marks in excluded and included exons, Chen et al. built a Bayesian network for included and excluded exons. In this work, they investigated the relationships between histone marks, so the direction of the specific marks was not assessed [44].

Curado et al. defined a small set of exons showing promoter-like signatures. These alternative exons show elevated inclusion with increased levels of H3K9ac, H3K4me3, and H3K27ac [10].

Studying AS in mouse brain after cocaine stimulation, Hu et al. found H3K36me3 and H3K4me1 positively associated with SE inclusion [47]. They also emphasize the importance of neighboring regions in predicting alternative event usage. In their more recent work, they studied SE in the development of the mouse brain and came to similar conclusions [48].

H3K79me2 is enriched at splice sites of SE and can reveal events specific to cancer types [49].

Xu et al. constructed *DeepCode*, a neural network based on sequence and epigenetic predictors, to predict exon inclusion-related events relevant to embryonic stem cell (ESC) fate decisions. Despite the limited interpretability of single features, H3K36me3 and H3K79me1 could be identified as the most important without a clear direction, except for specific genes [50]. In a later study, they focused on a comprehensive evaluation of MX and SE involved in ESC fate decisions [51]. In this context, they emphasized the diverging roles of H3K36me3 depending on the exon while showing a positive correlation with inclusion globally. This overall positive association was also found for H3K27ac and H4K8ac.

In another effort to combine sequence information with epigenetic features, Lee et al. built a recurrent deep learning model to predict SE inclusion, achieving an area under the ROC curve of 0.94 [52]. Globally, they emphasize finding a negative association for H3K27me3 and H3K9me3 to exon inclusion, while open chromatin, H3K27ac, H3K36me3, and H3K4me1 were positively linked.

Agirre et al. found splicing-associated chromatin signatures of epigenetic marks enriched in a subset of SE, depending on their inclusion level [53]. They found that the role of marks highly depends on the combination with others and their position, i.e., near the 5’/3’ splice site or inside the exon body. Overall, seven signatures were significantly associated with alternative exon inclusion categories. Although the analysis is genome-wide, the combinations are found in at most 165 exons at once.

Petrova et al. investigated retained intron (RI)s comprehensively in immune cells by building machine learning models and assessing their variables’ importance [54]. Following intrinsic features, epigenetic markers were most predictive for intron inclusion. Specifically, variables related to nucleosome-free regions near the upstream splice site were associated with retention. In addition, they could show for their data that the enrichment of H3K36me3 in excluded introns was only due to nucleosome occupancy.

It is important to note that these studies are based on various organisms, biosamples, definitions of splicing and exon inclusion, and utilize multiple statistical models. The heterogeneity of the study designs and software tools involved can explain some of the inconsistencies in the literature.

These partially contradictory results motivated us to study the reference epigenomes from the IHEC EpiATLAS resource to systematically assess evidence for the role of epigenetic mechanisms in regulating AS.

## 2 Results and Discussion

In this study, we consider 405 epigenomes from the EpiATLAS project for which RNA-Seq, DNAm, and six histone marks are available (Figure 1). RNA-Seq transcript expression files from RSEM [55] serve as SUPPA2 [56] input to identify splicing events and compute their proportion spliced-in (PSI) values. The PSI quantifies a given event by representing the proportion of transcripts including this event. We only consider transcripts with support level 1 or 2 to remove potential noise and filter for SE and RI on autosomes that do not overlap with other events or share their splice sites across genes. After filtering, 6408 of 15038 and 1440 of 2746 events with quantified PSI values remained for SE and RI, respectively. Each event was categorized into high or low variability based on the PSI standard deviation to investigate differences in events that are more or less dynamic. To assess epigenomic associations to exon/intron inclusion, we aggregate the epigenetic signal in the alternative region and its adjacent areas. Additionally, we include the previously introduced maximum promoter and summed enhancer signal of H3K27ac [57, 58] (see Methods). Finally, we gather the gene expression value and intrinsic features, such as the regions’ widths, distance to their TSS and transcription end site (TES), and splice site entropy using MaxEntScan [59].

**Fig. 1:**
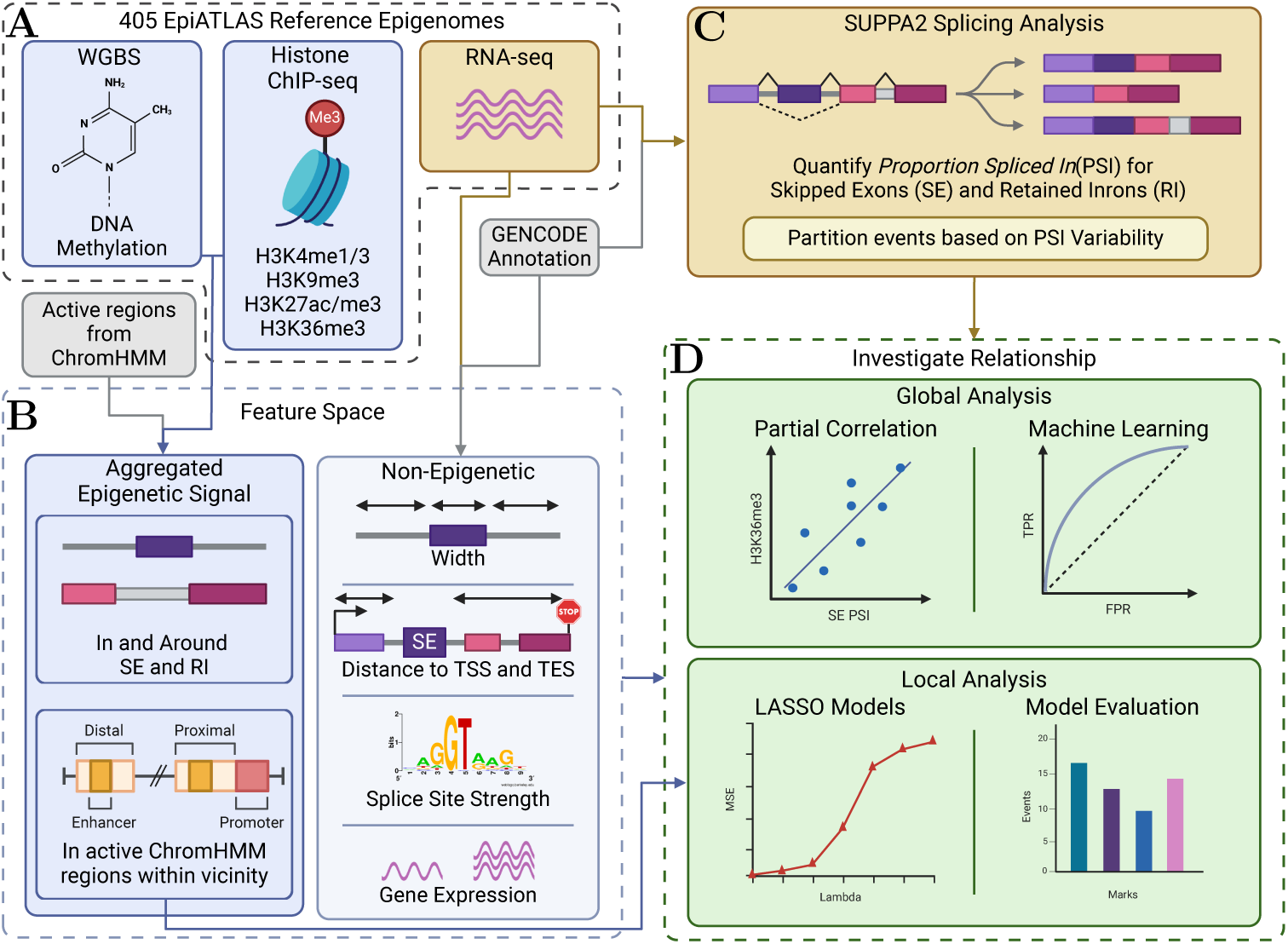
Overview of the study workflow. **A** Data acquisition from the EpiATLAS project, including whole-genome bisulfite sequencing (WGBS), histone ChIP-seq, and RNA-seq. **B** Preparation of epigenetic signals and integration with intrinsic features and gene expression data. **C** Detection and quantification of splicing events, focusing on skipped exons (SE) and retained introns (RI). **D** Analysis of epigenetic associations with alternative splicing using genome-wide partial correlation, machine learning models, and event-specific generalized linear models.

### 2.1 Revisiting genome-wide associations with splicing across diverse epigenomes

Many studies have reported global links between epigenetic marks and alternative splicing. We investigate genome-wide associations between histone modifications/DNA methylation and PSI values on the EpiATLAS dataset using partial correlation analysis and random forest machine learning models.

#### 2.1.1 Global correlation of epigenomic features weakens for variable events

The PSI values are inherently related to gene expression since they are computed using the same data. Thus, we calculate the partial correlation coefficient between the PSI and the explanatory features, controlling for gene expression. The genomewide partial correlation analysis shows that epigenetic features in and around the alternative regions are associated with alternative splicing besides intrinsic features such as splice site strength (Figure 2A). As expected, strong splice sites correlate positively with exon inclusion and negatively with intron retention. The widths of the alternative region follow the same trend. DNAm and H3K36me3 around the splice sites positively correlate with SEs’ PSI values, while H3K4me3 shows an opposing trend. In RI, H3K4me3 and H3K27ac are positively correlated with the intron PSI. DNAm in the downstream exon shows a negative correlation.

**Fig. 2:**
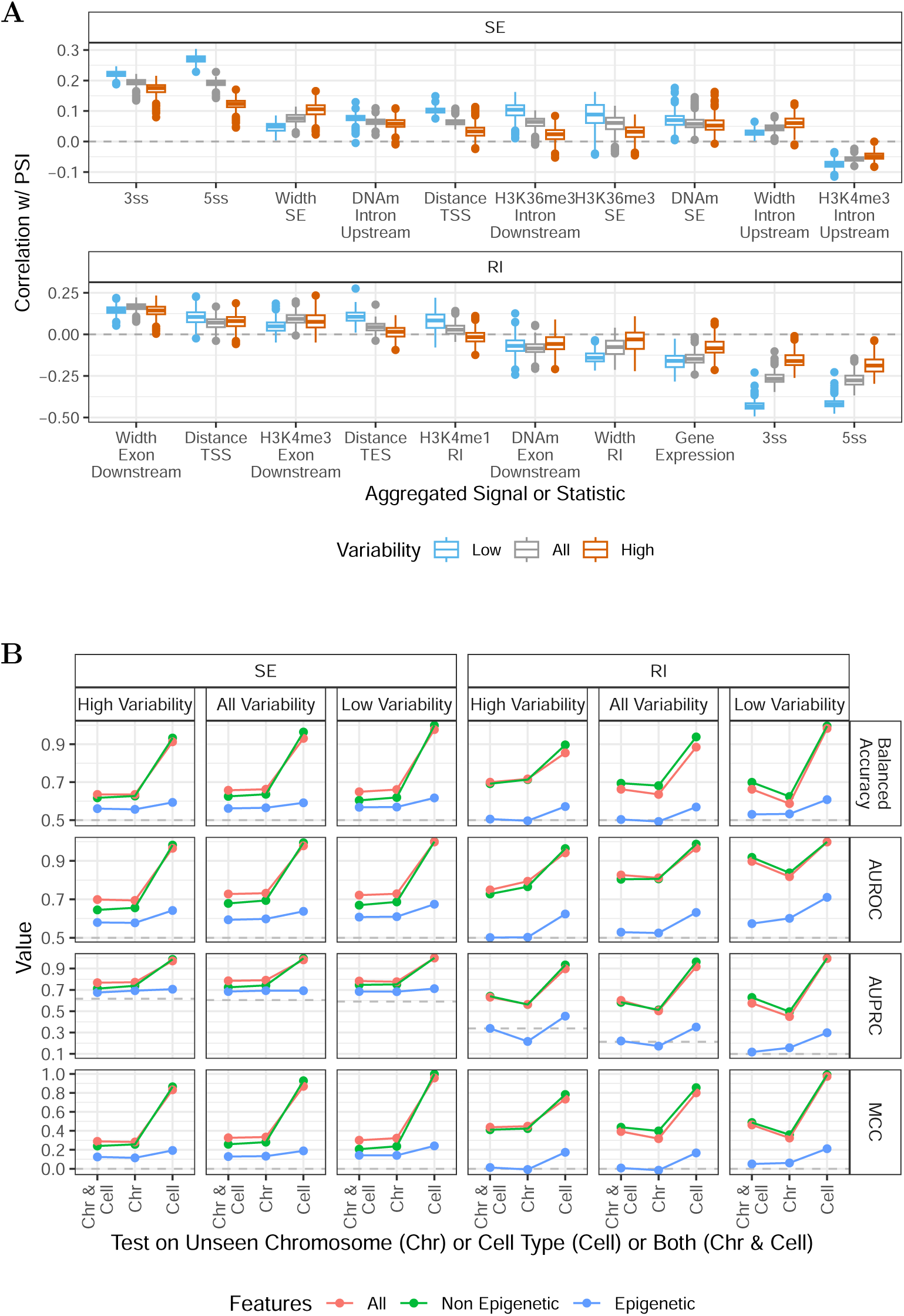
**A** Top 10 partial correlations betwe^9^en splicing event PSI values and epigenomic features, adjusted for gene expression. The analysis reveals a decrease in correlation strength for events with high variability compared to those with low variability. **B** Comparison of machine learning model performance in predicting exon/intron inclusion across different test sets. The models incorporating epigenetic features show a marginal improvement in accuracy compared to those using non-epigenetic features alone. Testing on unseen cell types improves performance compared to unseen events.

To investigate whether this correlation analysis merely reflects the underlying grammar of splicing without respect to variation, i.e., alternative splicing, we recomputed the partial correlation coefficients for each variability group. Figure 2A shows that the correlation changes depending on the variability. Specifically, most absolute correlations shrink for intrinsic and epigenetic features when only considering the dynamic events in the high variability group. This observation does not hold for the width of the exon and upstream intron in SE, which show opposite directions.

#### 2.1.2 Positive correlation of H3K36me3 and DNAm with exon skipping are supported by EpiATLAS

We next compared previously found genome-wide associations between AS and the epigenome to the trends in our partial correlation analysis to see where previous findings can be corroborated (Table 1; partial correlation coefficients are shown in Figure S4). For SE, H3K36me3 and DNAm are consistently associated with inclusion, as reported in most studies, whereas H3K4me3 and H3K27ac show mixed results. In contrast to some reports, no associations were found for H3K9me3, H3K27me3, and H3K4me1. In RI, the associations are less clear. Our results for DNAm, H3K4me1, and H3K36me3 do not align with the literature. Moreover, we found positive associations of H3K4me3 and H3K27ac that had not been reported before. The discrepancy between the literature and our results could be due to the different organisms, cell types, and experimental designs used in the studies.

**Table 1:** Directed associations of epigenetic marks with alternative exon/intron inclusion according to the literature and on our EpiATLAS analysis. If a study has conflicting global results due to the position of the mark, the association in the alternative region is noted. When the direction of the association is not specified or found to be non-significant, it is not shown in the table.

| Epigenetic Mark | Genomic Association | Exon Inclusion |  | Intron Inclusion |  |
| --- | --- | --- | --- | --- | --- |
|  |  | Literature | EpiATLAS Analysis | Literature | EpiATLAS Analysis |
| H3K9me3 | (Constitutive) Heterochromatin | ↑ [43]<br>↓ [37, 52] | — | | — |
| H3K27me3 | Heterochromatin/Suppression | ↑ [43]<br>↓ [40, 45, 52] | — | | — |
| H3K4me3 | Active Promoters | ↑ [40]<br>↓ [38, 42] | ↘ | | ↑ |
| H3K4me1 | Enhancers | ↑ [47, 48, 52]<br>↓ [38] | — | ↑ [40] | — |
| H3K27ac | Active Enhancers | ↑ [51, 52]<br>↓ [45] | ↘ | | ↑ |
| H3K36me3 | Transcribed Genes | ↑ [35–43]<br>[47, 48, 51, 52]<br>↓ [45] | ↑ | ↑ [40] | — |
| DNAm | Promoter Suppression/Transcribed Genes | ↑ [40, 60] | ↑ | ↑ [40] | ↘ |

#### 2.1.3 Epigenetic marks marginally improve inclusion prediction

To better understand how robust associations of epigenetic and intrinsic features are across chromosomes or cell types, we construct random forest binary machine learning models classifying the exclusion 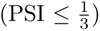 and inclusion 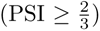 of exons/introns.

As a test set, we chose events from held-out chromosomes and cell types to avoid data leakage, namely chromosome 1 as well as B and T lymphocytes. Events with PSI values between 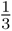 and 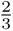 were discarded for this analysis, almost all of which were previously categorized as high variability. Additionally, we built models containing only or no epigenetic features to assess how much prediction accuracy is retained with a restricted feature space. We conducted this analysis again split by variability to assess potential differences in the prediction difficulty when considering more or less dynamic events. Figure 2B shows that, as expected from the correlation results, the epigenetic models without intrinsic features are worse at correctly recapitulating the classes. Interestingly, testing on seen events but unknown cell/tissue types leads to near-perfect performances when including intrinsic features, while testing unseen chromosomes with known cell types does not increase performance. This indicates that models cannot generalize well to new events, emphasizing distinct mechanisms per event over cell-type specificity.

Overall, RI performs better than SE compared to the respective random baseline and when excluding epigenetic features entirely, except for high variability introns (Figure S1B). Other than for low variability introns, RI models using only epigenetic information are not superior to random models when tested on unseen events. The ROC and PR curves in Figure S2 underscore this observation.

In addition to the random forest models, we evaluated generalized linear models for classification, namely LASSO and LASSO for feature selection, followed by logistic regression. Random forest performs better than the linear models, which could be due to their ability to pick up non-linear relationships (Figure S1A).

The variable importance of the random forest models recapitulates the findings from the partial correlation analysis (Figure S3). H3K36me3 and DNAm have the highest variable importance for the epigenetic SE models. Notably, for RI, H3K4me3 shows higher feature importance than DNAm. As previously noted, epigenetic features are not as important as intrinsic features, which is reflected in the better performance of the models leveraging both. Thus, the interpretation of the epigenetic model variable importance in genome-wide models is limited. Overall, removed events were mainly highly variable, leaving comparably more lowly variable events showing stronger links in Figure 2A. In contrast, the comparably better results on unseen cell or tissue type indicate that the locus-specific context of the splicing events is crucial for regulation, motivating a closer inspection of individual events.

### 2.2 Event-specific models predict PSI as well as on different chromosomes

Since a genome-wide approach does not capture event-specific trends, and intrinsic features cannot explain PSI changes for a single event, we construct a regularized linear model for each event modeling the PSI values based on the epigenetic neighborhood as well as close-by active elements defined by active ChromHMM states [61, 62]. These models identify 206 SE and 60 RI events with epigenetic features predictive of their PSI value and do not show a gene expression bias Figure 3A. Models with expression bias were defined as those that fit the PSI significantly better when gene expression was added to the epigenetic features as determined by an ANOVA test. We conduct these tests to assess whether the epigenetic marks are merely predictive of gene expression and can predict PSI through this proxy. Such models constitute 235 and 65 for SE and RI. Consequently, for 5506 and 1231 events, no features were selected by the LASSO models, indicating that these events are not epigenetically regulated.

**Fig. 3:**
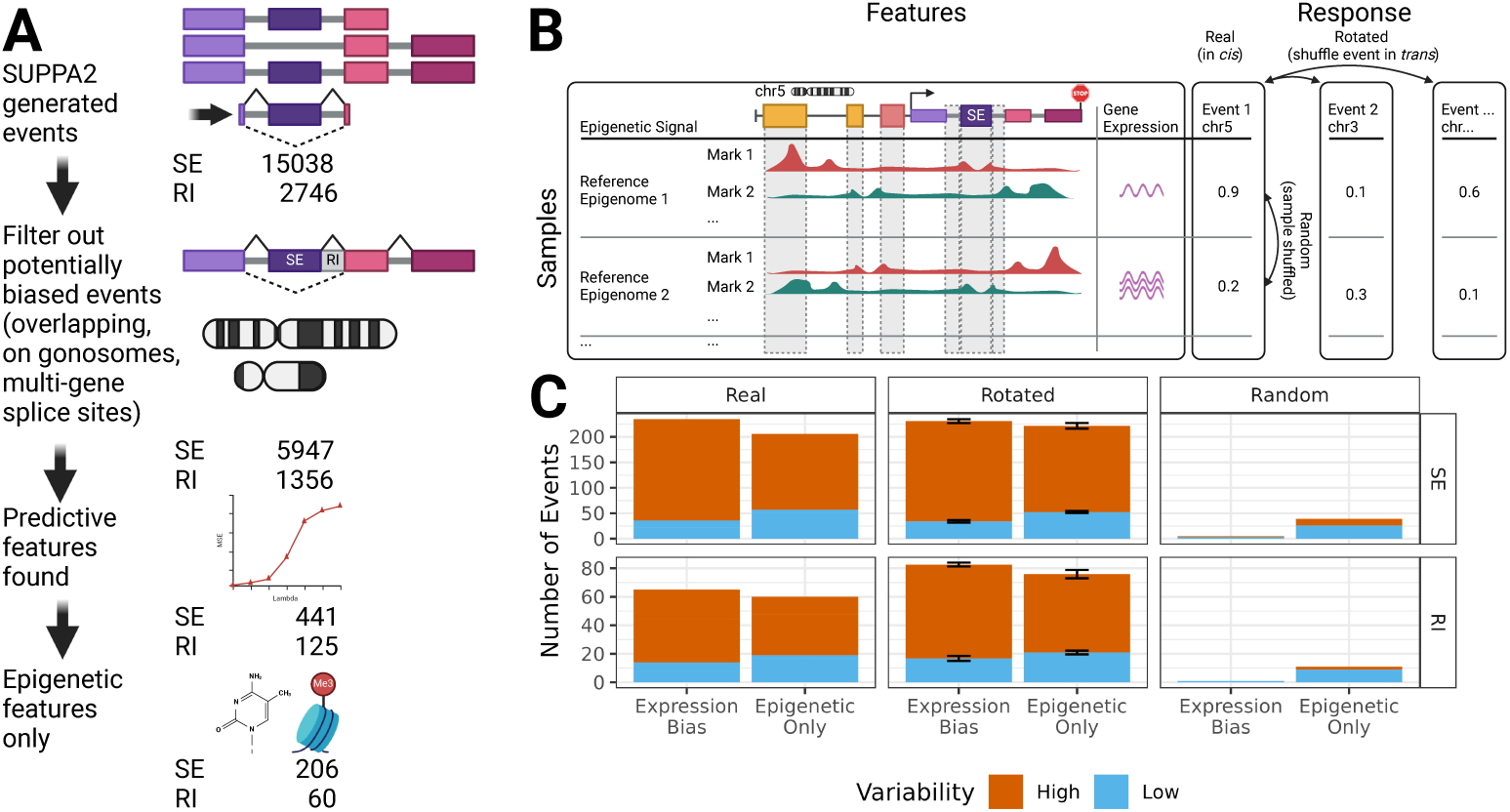
Evaluation of event-specific models. **A** The workflow for filtering potentially biased splicing events and identifying predictive epigenetic features. **B** Comparison background models using shuffled (random and rotated) PSI vectors. **C** Number of models built on real, random, and rotated responses, illustrating the impact of sample-specific epigenomes on splicing prediction.

Further, we build models on two types of shuffled response vectors to assess how often predictive features would be found by chance (Figure 3B). Firstly, random responses are created by using the original PSI vector and shuffling it such that the values still belong to the same event but a different sample. Rotated responses represent the PSI vector from a different chromosome, such that the event is different, but responses are matched to the original sample. This way, for each feature matrix, we can evaluate whether a model picks up potential cis-regulatory epigenetic effects on the splicing outcome or simply a sample-specific fingerprint.

Figure 3C shows that random models rarely find predictive features, while rotated models are created for a comparable number of events. The distribution of high and low variability events appears alike for both real and rotated models. This similarity supports the hypothesis that sample-specific epigenomes can predict sample-specific PSI values. Hence, it is unclear if the features we find in cis are mechanistically linked to splicing.

#### 2.2.1 Features between real and rotated models differ but not in characteristics

To explore if the models systematically prefer specific epigenetic marks in explaining splicing outcomes, we count how often these have been selected across models without expression bias (Figure 4A). Note that we count the use of each epigenetic mark only once, even though models could select multiple epigenetic features from the same mark. Although the distribution differs for the randomized responses, real and rotated marks look similar. According to the global analysis and the literature, H3K36me3 should predict splicing outcomes, but the models do not preferentially select this histone modification. Instead, especially DNAm is selected more often than other marks.

**Fig. 4:**
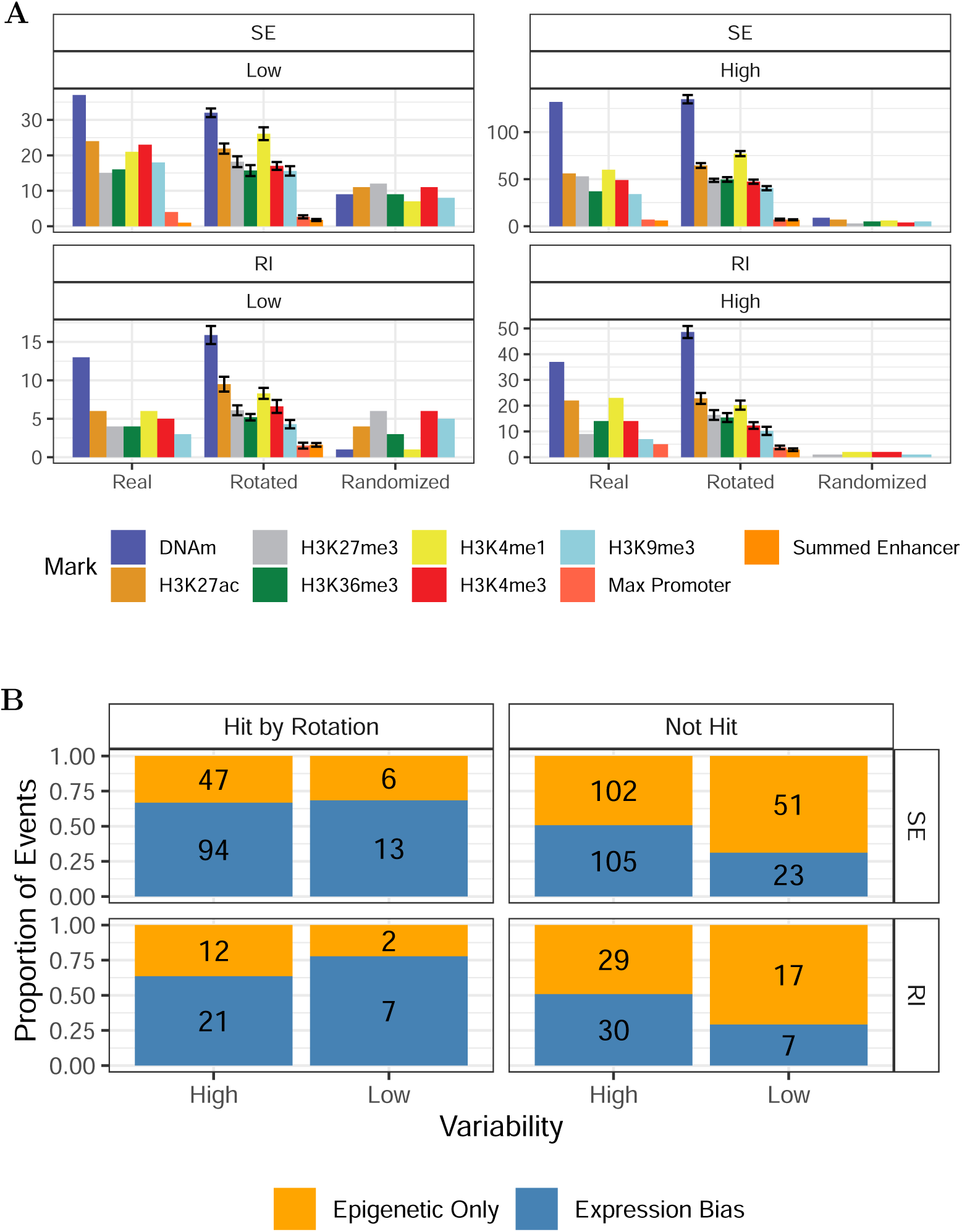
**A** Distribution of predictive epigenetic marks in models without expression bias, showing a preference for DNAm across both real and rotated models, potentially indicating sample-specific effects. **B** Comparison of the proportion of epigenetic-only and expression-biased models when separating splicing events that models with rotated PSI values could predict.

Since DNAm can be highly cell-type and sample-specific, this observation aligns with the hypothesis that the selected features for both real and rotated models represent proxies for cell type instead of offering potentially mechanistic explanations. Another explanation could be that DNAm offers a cleaner signal than the comparably noisy ChIP-seq data.

Further, we investigated which events can be predicted with features from other chromosomes. Following the rotation procedure outlined in Figure 3B, we checked whether there are models that select *trans* features, i.e., from the feature table from an event on a different chromosome, to predict an event’s splicing outcome (PSI vector). Those events predicted in *trans*, which we call ”hit” by the rotation background (Figure 4B), make up less than half of the events for which models could be built. Interestingly, these events more often show expression bias, as opposed to the ones not ”hit,” which have a higher proportion of epigenetic-only models. This observation suggests that epigenetic features on a different chromosome help predict across-chromosome gene expression patterns and, thus, in some cases, splicing.

Consequently, we investigated whether features predictive in *trans* are also important for predictions in *cis*. To this end, we gathered all features with nonzero coefficients in the rotated, across-chromosome LASSO models per event and fit the real response again without those features. For SE and RI, respectively, 359 and 85 models also find predictive features on the filtered feature space, while 82 and 40 do not select any epigenetic variables. Interestingly, 14 and 3 models select features only on the filtered features, likely due to the smaller feature set’s effect on the lambda parameter selection that governs the regularization in LASSO. Of the events for which models were built for both feature spaces, the chosen variables do not change for 65.7 and 62.4 percent, indicating more confidence in their robustness.

#### 2.2.2 No clear preference between long-range and local epigenetic signals for splicing prediction

Initially, we hypothesized that regions at a mid-range distance to the event are most relevant, so we selected 50kb as the window size. To investigate whether long-range regions hold more information or whether local epigenetic signals are sufficient, we conducted experiments by extending (windows size of 500 kb) and restricting the feature space. For the latter, we considered only adjacent epigenetic signals without including active regions from the ChromHMM annotation. Figure S5 shows that an increased number of explanatory variables helps build more models in which predictive features are retained. Analogously, restriction to the local signal leads to approximately half the number of models compared to the original feature space. However, the number of low variability models without expression bias approximates the random response. Still, the relationship to the rotated models, i.e., a similar number of models for real and rotated responses, remains for all three ways of defining the feature space. Comparable observations can be made on the level of selected marks and the analysis of responses that could be predicted from another chromosome (Figure S6). However, low numbers, especially for local models, do not reflect all trends. The differences in the results may also be due to differences in model complexity, i.e., more predictive models can be found if more features are available for training. Adjusting the window size thus did not clearly show whether the long-range or local signal is most important for the splicing outcome.

#### 2.2.3 Previously identified epigenetic regulation is not captured by the models

We have used the EpiATLAS dataset to construct event-specific models for predicting PSI values based on epigenetic features and gene expression. From the 14 exons previously reported in the literature to be regulated by splicing, only three models were among those with predictive epigenetic features (Table 2). Those three models use the extended feature space, including active regions around the splice sites in a 500 kb window. If exons were excluded due to overlap or multi-gene splice sites, we included them for this analysis but did not change the TSL filter.

**Table 2:**
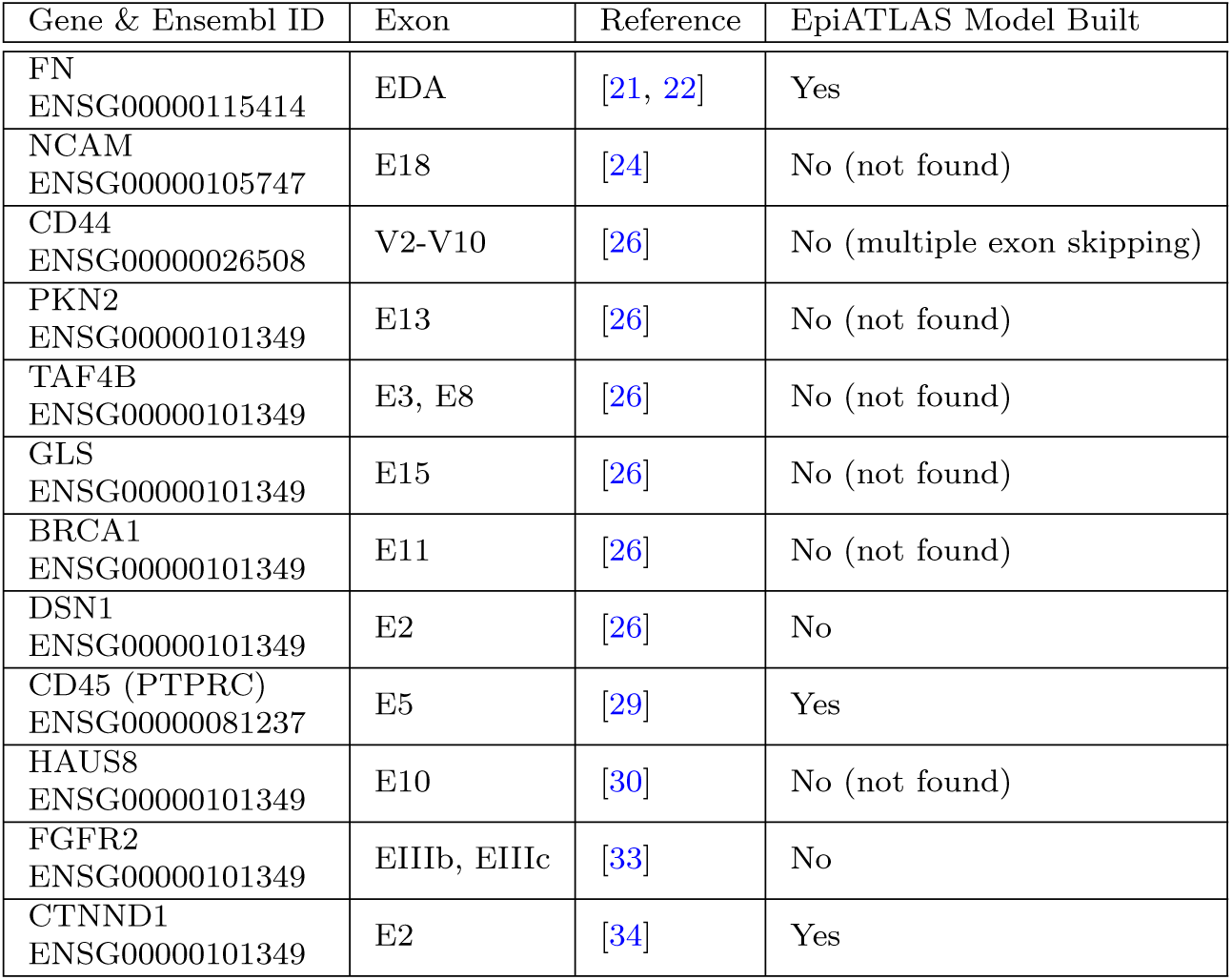
Previously identified exons regulated by epigenetic marks, with an indication of whether predictive models were successfully constructed using the EpiATLAS dataset. The table underscores the challenges of replicating known regulatory mechanisms in the context of alternative splicing.

| Gene & Ensembl ID | Exon | Reference | EpiATLAS Model Built |
| --- | --- | --- | --- |
| FN<br>ENSG00000115414 | EDA | [21, 22] | Yes |
| NCAM<br>ENSG00000105747 | E18 | [24] | No (not found) |
| CD44<br>ENSG00000026508 | V2-V10 | [26] | No (multiple exon skipping) |
| PKN2<br>ENSG00000101349 | E13 | [26] | No (not found) |
| TAF4B<br>ENSG00000101349 | E3, E8 | [26] | No (not found) |
| GLS<br>ENSG00000101349 | E15 | [26] | No (not found) |
| BRCA1<br>ENSG00000101349 | E11 | [26] | No (not found) |
| DSN1<br>ENSG00000101349 | E2 | [26] | No |
| CD45 (PTPRC)<br>ENSG00000081237 | E5 | [29] | Yes |
| HAUS8<br>ENSG00000101349 | E10 | [30] | No (not found) |
| FGFR2<br>ENSG00000101349 | EIIIb, EIIIc | [33] | No |
| CTNND1<br>ENSG00000101349 | E2 | [34] | Yes |

The skipped exon in the CD45 gene [29] could be identified, and the regularization models selected the DNA methylation in two active regions as predictive for the PSI value. Those two regions appear to be in different TADs and seem far away; one overlaps with another gene (Figure S7A). Thus, these are likely spuriously correlated features.

We identify the EDA exon in the FN1 gene [21, 22] for which H3K4me1 and DNAm in active regions predict the PSI value. Figure S7B shows that both regions overlap with other genes and are unlikely to control the alternative splicing of the exon in question in *cis*.

Lastly, a CTNND1 exon [34] has predictive features selected. Still, 25 epigenetic features are found in 89 samples. Because of the low sample-to-feature ratio, this model should not be considered reliable.

## 3 Conclusion and Future Perspectives

Many studies have previously reported associations between epigenomic marks and changes in AS. However, some of these studies had conflicting results, which motivated us to systematically reassess the evidence for epigenetic control of alternative splicing leveraging the comprehensive EpiATLAS data set. In line with previous studies, our global analysis successfully recovered known associations, particularly the positive correlation between DNA methylation (DNAm) and H3K36me3 with exon inclusion. However, our study is the first to report that these associations diminish when focusing on highly variable events. This finding underscores the critical role of variability for the epigenetic associations to alternative splicing while indicating other mechanisms might be determinants for dynamic regulation. Beyond a global association analysis, the large sample size of the EpiATLAS enabled us, for the first time, to model splicing outcomes from epigenetic data on an event-specific basis. Still, we could not reproduce all exon-specific results from the literature, which shows the problem of replicating single studies on atlas-scale resources. Moreover, the results of our analysis emphasize that the relationship between the epigenome and splicing outcomes is specific to individual cell types. This specificity complicates disentangling the epigenetic effects on splicing, even within such a comprehensive and uniformly processed dataset.

Our study has several limitations. First, assuming linearity in our event-specific models likely oversimplifies the intricate and often nonlinear nature of splicing regulation. This simplification may not accurately capture the complex regulatory mechanisms at play but is deemed acceptable due to the limited number of samples and the benefit of more straightforward interpretability. Though effective in highlighting differences, the straightforward definition of variable events may not fully encompass the multifaceted nature of splicing variability. Further, the selection of histone marks provided by IHEC may not include those most relevant to alternative splicing regulation but appear more closely linked with regulating gene expression and cell type [63]. Moreover, the absence of data on nucleosome occupancy and other potentially significant epigenetic marks, like H3K9ac, H2BK12ac [38, 45], or H4K20me1 [41], limits the scope of our findings.

Future research should focus on enhancing computational models by integrating information on splicing factor activity, which could provide a more comprehensive understanding of the regulatory landscape. For instance, combining SF expression with sequence context and epigenetic signal from DNAm to nucleosome occupancy and histone marks could describe the splicing regulatory landscape more accurately. Such studies should begin on specific cell types to avoid the confounding effects of mixed cell types. Single-cell multi-omics technologies and long-read RNA sequencing promise a more accurate and detailed epigenomic and splicing profile to train and validate such computational models. Furthermore, perturbation experiments, such as CRISPR screens, will be essential in validating epigenetic influences on alternative splicing [34].

In conclusion, our study highlights the cell-type specificity of the interactions between epigenetics and splicing outcomes on a large, high-quality dataset coupled with rigorous methodological controls. However, we acknowledge the limitations of our approach and propose several avenues for future research that could further elucidate the complex relationships between epigenetic modifications and splicing outcomes. The broader implications of our findings suggest that careful in silico testing should be standard practice in epigenomics, given the complexity and variability inherent in these data.

## 4 Methods

If not indicated otherwise analyses were conducted using R version 4.2.1 [64], GenomicRanges version 1.48.0 [65], ggplot2 version 3.4.4 [66], and data.table 1.15.4 [67]. The code is available at https://github.com/daisybio/IHEC-AS under the GPL-3.0 license.

### 4.1 Prepared Datasets

All analyzed data was derived from the EpiATLAS project and is available through the IHEC data portal (https://epigenomesportal.ca/ihec/) [19, 20]. Data accessions and associated public metadata are available through EpiRR (https://www.ebi.ac.uk/ epirr). 405 reference epigenomes with observed RNA-seq and WGBS data and either observed or imputed [68] histone ChIP-seq for all six histone marks were identified and downloaded via SFTP. Methylation data was extracted from per chromosome CpG matrices containing the percentage of methylated reads at that position with a minimum coverage of 10. Histone mark signal was extracted from observed or imputed MACS2 [69] p-value signal bigWig files. GENCODE v29 [70] was used as genome annotation and filtered for transcripts with transcript support level (TSL) one or two. Isoform result tables from RSEM [55] are used and transcripts per million transcripts (TPM) values are computed on all TSL1/2 transcripts using expected counts and effective lengths. SUPPA2 version 2.3 [56] identified and quantified AS events. We ran the tool on all available event types and configured it to pool genes with shared splice sites across genes for event identification and a TPM filter of one for quantification. SE and RI events were filtered based on their characteristics; events not satisfying at least one requirement were discarded:

- The event is on an autosome, not on a gonosome or scaffold
- The event does not share splice sites across genes
- The alternative region of the event does not overlap with the alternative region of another event of the same type.
- The alternative region or the adjacent regions of the event do not overlap with the alternative region or the adjacent regions of another event of another type.

We computed the standard deviation (SD) for each event across the reference epigenomes to assess the variability of single events. To combat inflated SD values for events with high *NA* proportions, we always divide by the total number of samples, i.e., 405 *−* 1.

### 4.2 Genome-wide associations

#### 4.2.1 Feature definition

We first gather intrinsic features for the global partial correlation analysis and prediction models. The width of the alternative region and its upstream and downstream adjacent regions are directly inferred from the event identifier provided by SUPPA2. We compute the distance to the TSS and TES by taking the most upstream or downstream TSS or TES of all transcripts, including the region of interest, and computing their distance to the alternative region. Widths and distances are *log*2 transformed. Splice site scores are computed for the alternative region, i.e., for SE, the 5’ site refers to the downstream site, while for RI, it is the upstream site. Sequences of length 9 for 5’ and 23 for 3’ sites were extracted from the reference genome assembly GCA 000001405.15 with the Bioconductor Biostrings package [71] and scored via MaxEntScan [59] using the scripts available on their website with Perl version 5.32.1. The overall gene expression is defined as the *log*2 sum of all the transcript TPM values belonging to that gene.

We gather the mean epigenetic signal in the alternative and adjacent regions. For the histone marks, we utilized bigWigAverageOverBed [72].

Additionally, we included the H3K27ac maximum promoter value, which represents the highest H3K27ac signal detected within *±*200 bp around any annotated TSS of a gene. We also incorporated the summed enhancer signal for each gene, calculated by aggregating the H3K27ac signal across all associated enhancers [58]. These enhancers were identified based on the predicted interactions using the gABC model, which assigns enhancer activity in proportion to the frequency of contacts with the gene’s promoter [57]. The corresponding files (GeneMaxPromoterSignal.txt.gz, gABC GeneSummedEnhancer.txt.gz) were downloaded from the FTP server.

#### 4.2.2 Partial Correlation Analysis

Pearson’s partial correlation between each feature and the PSI value correcting for gene expression was computed for every reference epigenome using the ppcor package version 1.1 [73]. For gene expression, this corresponds to the standard Pearson correlation coefficient.

#### 4.2.3 Machine Learning Models

Each event-reference epigenome combination serves as a single sample in the prediction models. In cases where *NA* values occur, we replace them by the mean of the feature. All predictors are scaled to zero mean and unit variance before training. The corresponding test sets are scaled with the respective values from the training set. Their PSI values were categorized as excluded or included based on thresholds of less than 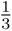 and greater than 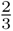, respectively. For a random baseline, all models are also trained on shuffled labels. Random forest classifiers were trained using the ranger package version 0.16.0 [74]. The caret package version 6.0.94 [75] assisted in data splitting, cross-validation, and hyperparameter selection. We held out events on chromosome one and from reference epigenomes from T lymphocytes or B lymphocytes as a test set. We conducted ten-fold cross-validation on the remaining data set with grouped splits by chromosome. We selected the configuration of hyperparameters with the lowest model complexity within one standard error of the best-performing configuration according to balanced accuracy. The ModelMetrics package version 1.2.2.2 [76] computed the Matthews correlation coefficient (MCC) and the PRROC package version 1.3.1 [77, 78] computed the receiver operating characteristic (ROC) and precision-recall curves and the corresponding areas under the curve.

### 4.3 Event-specific models

In addition to the gene expression and the epigenetic features mentioned above, the event-specific models also consider the epigenetic signal in potentially active regions as identified by ChromHMM [61, 62] in the vicinity of the alternative splicing event. The regions were downloaded from the IHEC FTP server (Stacked-ChromHMM V1 EnhancerStates.bed.gz). We consider a window of 50kb upstream and downstream from the alternative region. In addition, we also build models based on a larger window (500kb) and purely local models without active regions. The continuous PSI value serves as the response variable for the event-specific models. We do not consider events without variability in their target or with less than 25 samples. Thus, we build models for 5947 SE and 1356 RI events. Models can not be built if fewer groups are available than the number of folds (5) or if the response does not vary in a validation set of a given grouped fold.

Firstly, we train a LASSO model on a single event using the glmnet package version 4.1.8 [79] with five-fold cross-validation grouped by ontology to determine the lambda parameter, which is selected to be the highest value within one standard error of the best-performing lambda in terms of mean squared error (MSE). In this process, gene expression is without penalization to address potential biases, where the epigenetic signal would only predict overall expression. If nonzero variables other than gene expression remain, we check whether those have been selected in at least two cross-validation folds in addition to the whole model on the full dataset. We gather those features and fit an ordinary least squares (OLS) model using the epigenetic features and another one with epigenetic features and gene expression. Later, we can test whether adding gene expression has significantly enhanced the model’s ability to fit the PSI value.

For each feature table, we repeat this process on a randomized response vector. For the rotated models, we select another event’s response where all the PSI values of the current event’s reference epigenomes are also available, that event lies on a different chromosome, and is in the same variability group (i.e., low or high) as the current event. We repeat the process mentioned above for ten distinct rotated responses.

## Appendix A Supplementary Figures

**Fig. S1:**
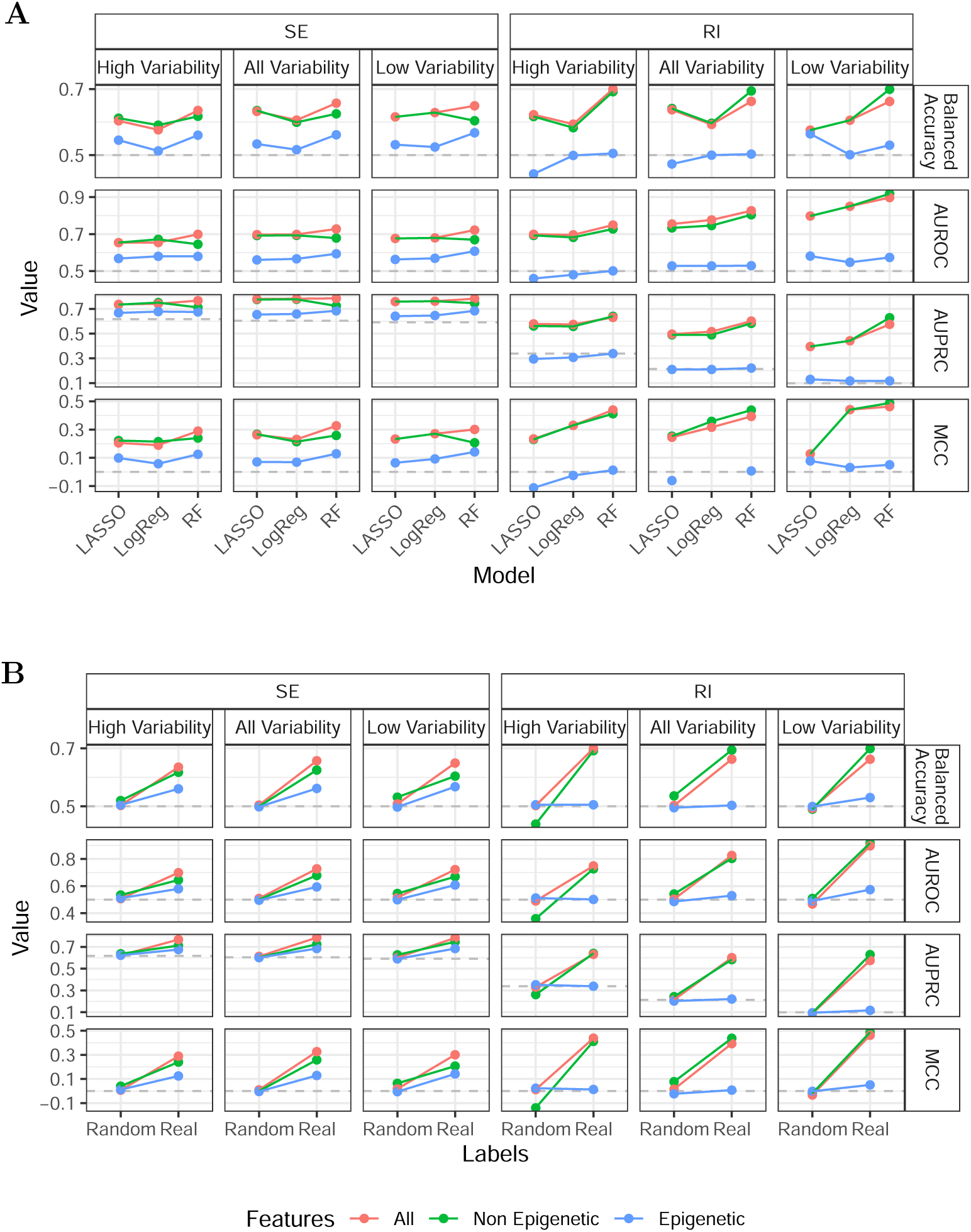
**A** Performance comparison of LASSO, logistic regression, and random forest models in predicting splicing events across different levels of variability. **B** Performance comparison of random forest models with a random baseline, i.e. shuffled labels, in 23 predicting splicing events across different levels of variability.

**Fig. S2:**
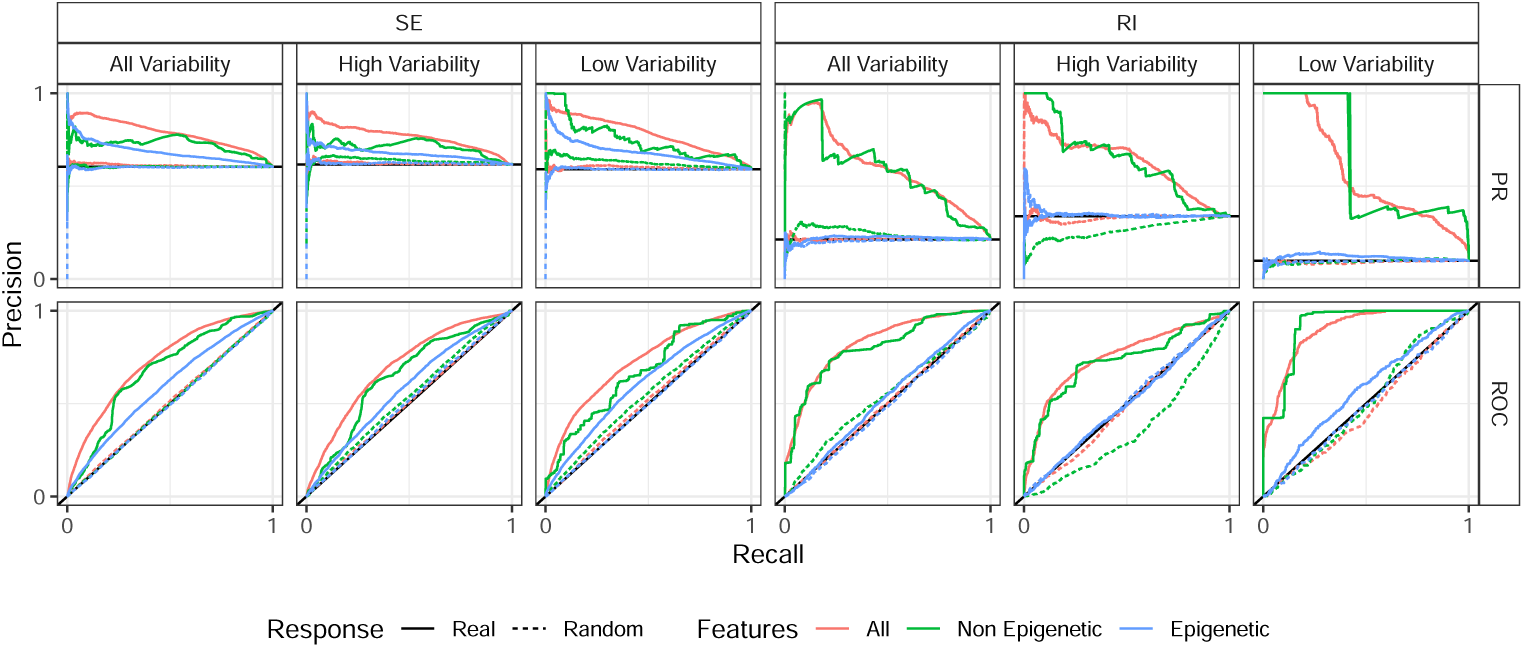
Receiver operating characteristic (ROC) and precision-recall (PR) curves for machine learning models predicting exon/intron inclusion, evaluated on real and randomized datasets. The curves illustrate the predictive power and reliability of models incorporating epigenetic features.

**Fig. S3:**
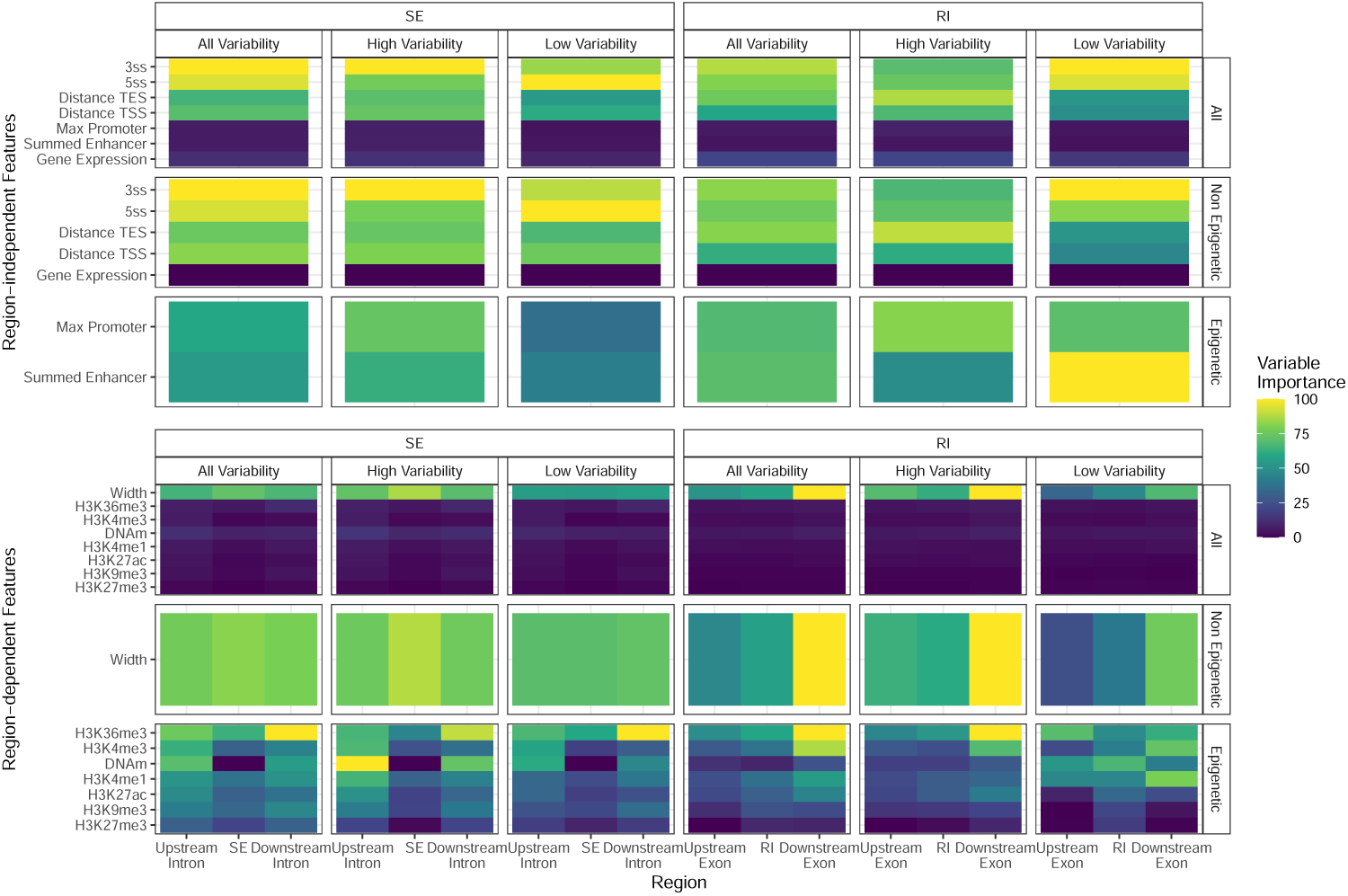
Variable importance of random forest model features for predicting splicing events. The figure shows the relative importance per model.

**Fig. S4:**
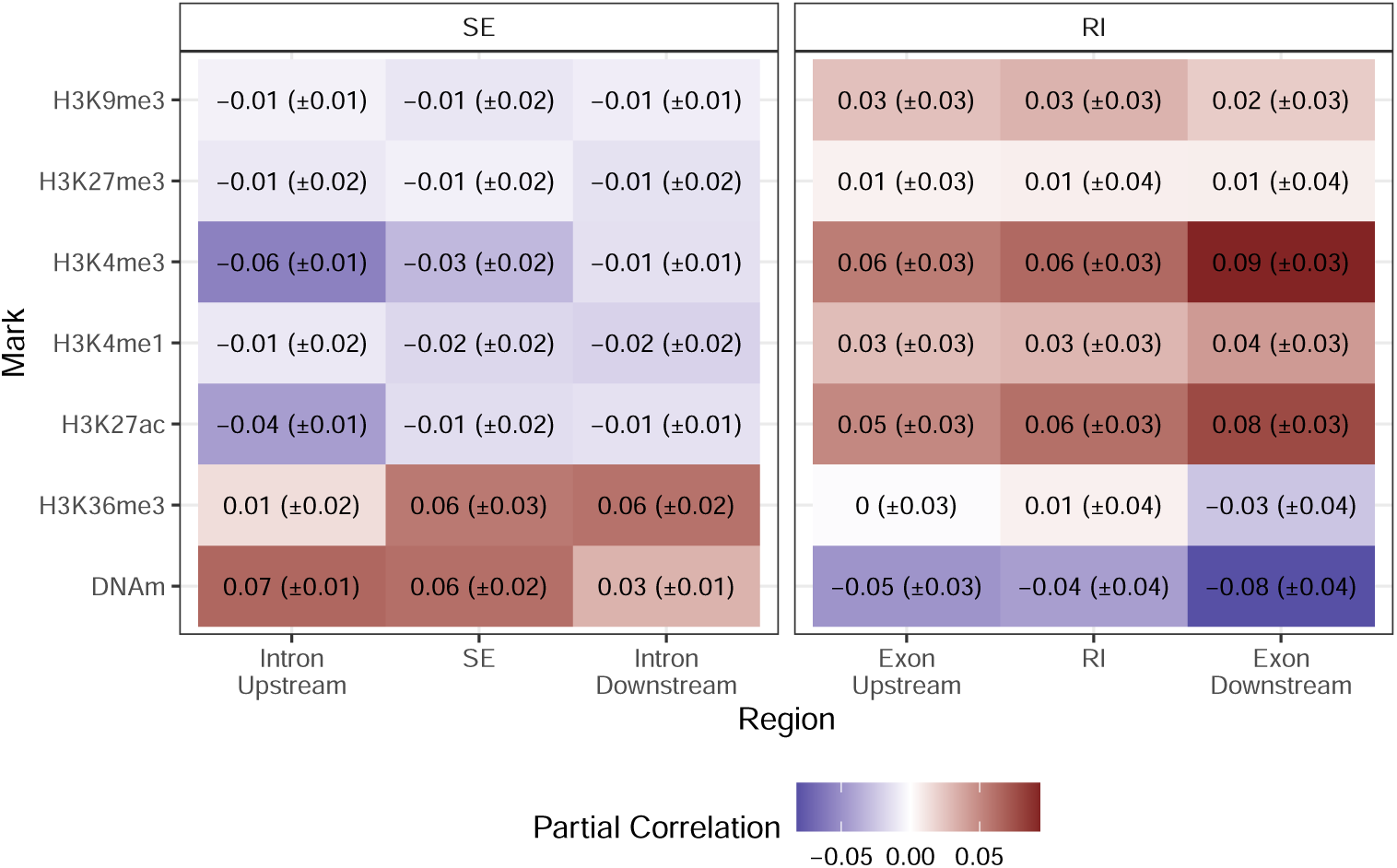
Mean (SD) partial correlation values between all splicing event PSI values and epigenomic features, controlling for gene expression.

**Fig. S5:**
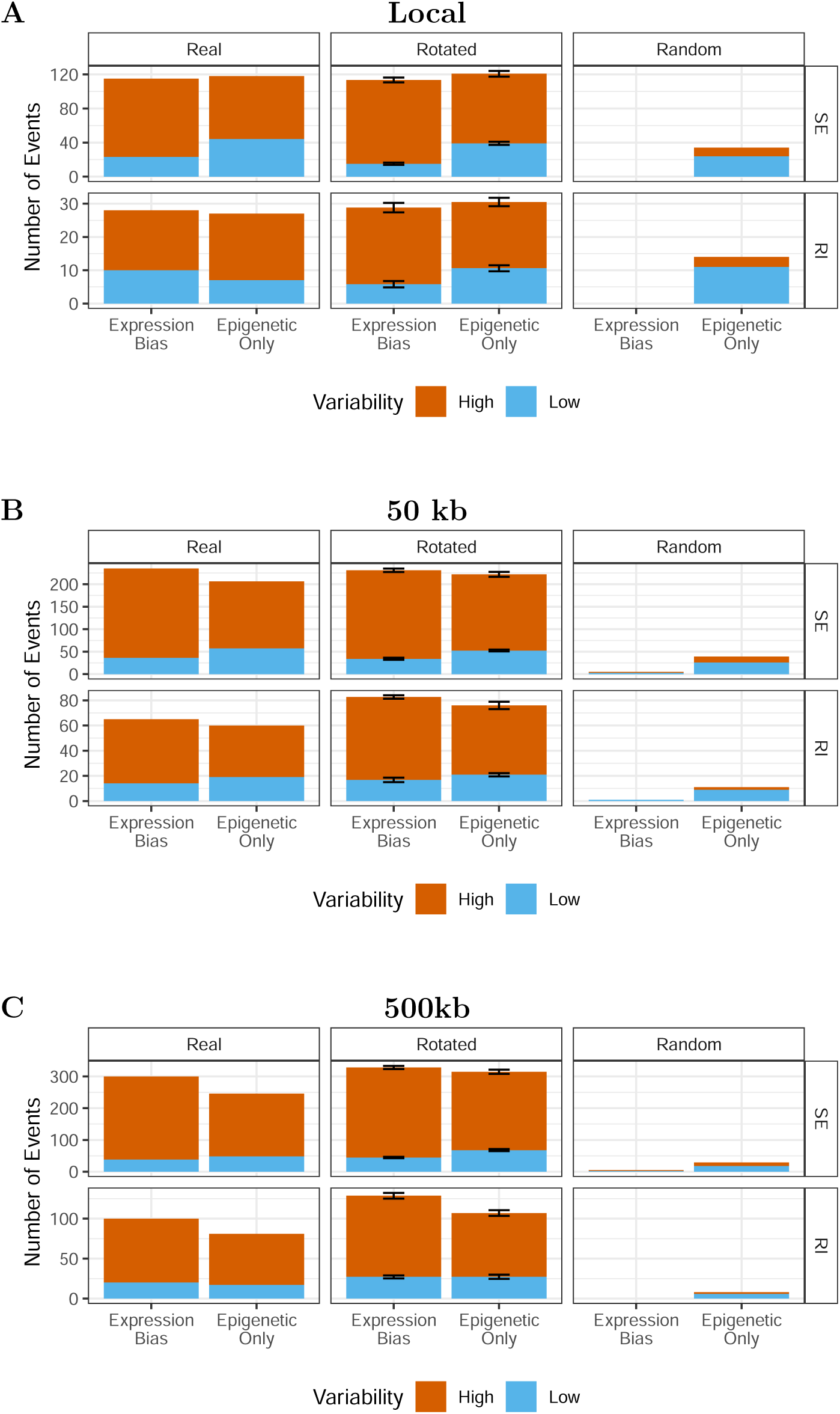
Impact of genomic distance on the identification of predictive epigenetic features for splicing events. **A** Local **B** 50 kb **C** 500kb contexts are examined to assess how the inclusion of neighboring regions influences models for predicting PSI values. The analysis compares real, randomized, and rotated data to evaluate the robustness of the predictive features identified.

**Fig. S6:**
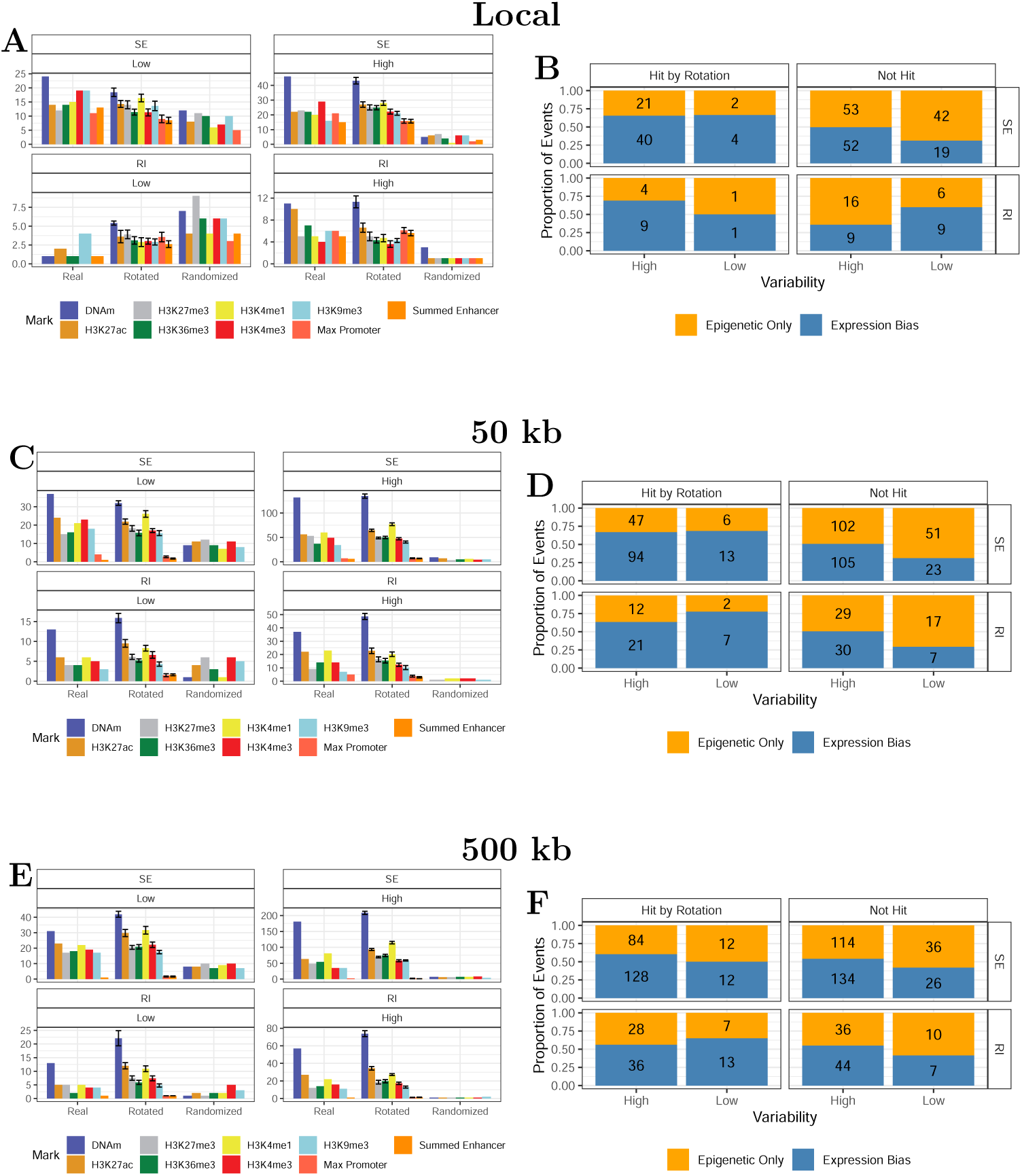
Impact of genomic distance on the predictive features for splicing events. **A, B** Local **C, D** 50 kb **E, F** 500kb contexts are examined to assess how the inclusion of neighboring regions influences predictive features and the comparison between epigenetic only and expression bias models respective to whether they have been predicted with across chromosome features.

**Fig. S7:**
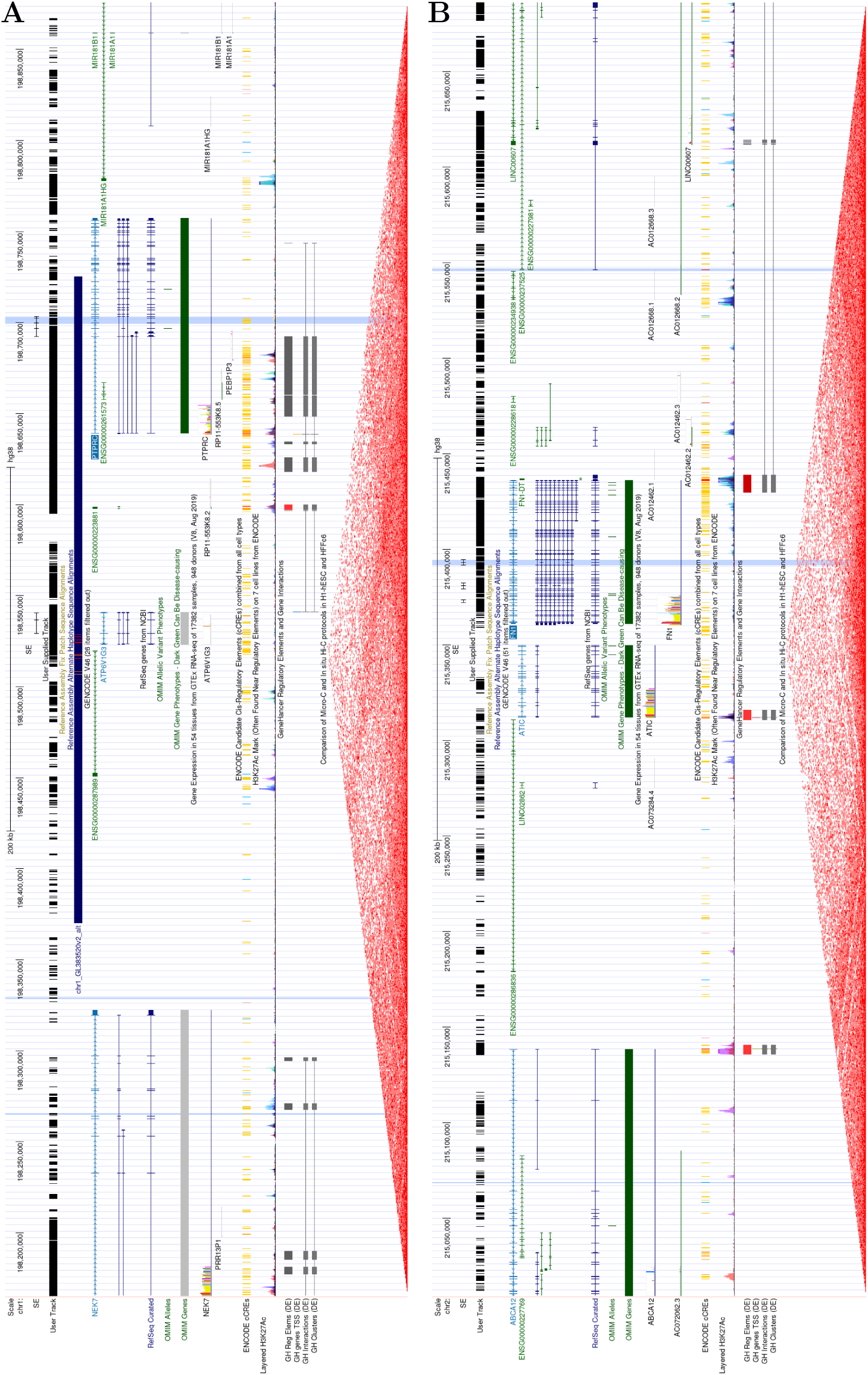
Screenshots of the genome browser with the predictive regions and the SE.

## Appendix B List of Abbreviations

AS: alternative splicing
ChIP-seq: chromatin immunoprecipitation sequencing
DNA: deoxyribonucleic acid
DNAm: DNA methylation
HP1: heterochromatin protein 1
IHEC: International Human Epigenome Consortium
MCC: Matthews correlation coefficient
MX: mutually exclusive exons
mRNA: messenger RNA
MSE: mean squared error
OLS: ordinary least squares
Pol II: RNA polymerase II
PSI: proportion spliced-in
RI: retained intron
RNA: ribonucleic acid
ROC: receiver operating characteristic
SD: standard deviation
SE: skipped exon
snRNP: small nuclear ribonucleoprotein
SRSF1: serine/arginine-rich splicing factor 1
TES: transcription end site
TPM: transcripts per million transcripts
TSL: transcript support level
TSS: transcription start site

